# Translating Ethiopian potato seed networks: identifying strategic intervention points for managing bacterial wilt and other diseases

**DOI:** 10.1101/2024.02.12.579952

**Authors:** Berea A. Etherton, Aaron I. Plex Sulá, Romaric A. Mouafo-Tchinda, Rogers Kakuhenzire, Haileab A. Kassaye, Frezer Asfaw, Vasilios S. Kosmakos, Richard W. McCoy, Yanru Xing, Jiahe Yao, Kalpana Sharma, Karen A. Garrett

## Abstract

**Context:** Informal seed trade can exacerbate crop disease epidemics. Potato production across Ethiopia is threatened by the spread of seedborne pests and pathogens, particularly by bacterial wilt, caused by the *Ralstonia solanacearum* Species Complex (RSSC). The RSSC is commonly dispersed through informal trade of seed potato, with the potential to move long distances across Ethiopia and among trading countries. Efficient disease testing programs and formal seed systems can reduce the risk of disease expansion in a country’s potato cropping system.

**Objective:** In this study, we characterized networks of potato value chain actors. We also identified candidate locations for disease surveillance and management interventions for improved seed systems, and locations at high risk for bacterial wilt establishment. We propose strategies to reduce the spread of bacterial wilt via infected seed.

**Methods:** We surveyed seed potato stakeholders to characterize interaction networks of potato value chain actors with a special focus on stakeholders engaged in seed potato quality assurance. We collected data regarding Ethiopia’s potato seed systems and analyzed the risk of spread of RSSC and other pathogens across the country through expert knowledge elicitation. Network metrics were used to characterize the informal seed trade system across Ethiopia, simulating pathogen spread across a network through scenario analyses. We characterized potato exports and imports to identify the risk of bacterial wilt movement through Ethiopia’s formal trading partners and neighboring countries where bacterial wilt has not yet been reported.

**Results:** Ware potato farmers and traders were reported to have weak communication with other stakeholders in the potato value chain. In network analyses and simulated epidemics, locations in Agew Awi, Gamo, Gofa, Kembata and Tembaro zones were identified as candidate priorities for national surveillance of pathogen invasion and expansion through management interventions and formal seed system development. Ethiopia has formal trade with Sudan, Southern Sudan, Russia, and other countries where bacterial wilt has not been reported. Ethiopia may be at risk of reintroduction of the RSSC from countries where it is present, like Kenya and India.

**Significance:** Improving seed systems to manage *R. solanacearum* and other seedborne pathogens is important for supporting food security and the livelihoods of smallholder farmers in Ethiopia. Implementing surveillance systems and management programs in locations like those identified in Agew Awi, Gamo, Gofa, Kembata, and Tembaro zones, and improving the communication between ware potato traders and other stakeholders, can help to strengthen informal trade of seed potato and mitigate bacterial wilt spread in infected seed.

## 1.0 Introduction

Food systems and food security are threatened by the spread of pathogens and pests, where effective stakeholder interactions and disease interventions in seed systems are crucial for managing disease outbreaks (Andrade-Piedra et al., 2022; Etherton et al., 2023; Ristaino et al., 2021; Thiele and Friedmann, 2020). Potato is a staple crop in many countries, including in sub-Saharan Africa, where it is currently threatened by a range of pests and pathogens (Kennedy et al., 2019; Petsakos et al., 2019; Tafesse et al., 2020). Ethiopian farms are at risk for disease spread and establishment, as systems for monitoring disease are uncommon and informal seed systems dominate the seed trade (Sperling et al., 2013; Sperling et al., 2020; Tessema and Seid, 2023; Thomas-Sharma et al., 2017; Thomas-Sharma et al., 2016). In this study, we consider potato seed systems in Ethiopia, focusing on stakeholder interactions and efficient locations for disease surveillance, and propose measures for improving these systems.

One important threat to potato in Ethiopia is bacterial wilt, caused by the *Ralstonia solanacearum* species complex (RSSC), which causes vascular wilt disease in other solanaceous crops (Abdurahman et al., 2017; Lemessa and Zeller, 2007). Bacterial wilt (also known as brown rot of potato) is caused by different races and biovars, however, *R*. *solanacearum* race 3 biovar 2 (or Phylotype II sequevar 1) is the primary strain associated with bacterial wilt of potato across Africa, where “race” distinguishes the plant host range and biovar distinguishes carbohydrate metabolism (Charkowski et al., 2020; Kurabachew and Ayana, 2017; Lowe-Power et al., 2020; Vailleau and Genin, 2023; Yaynu and Korobko, 1990). *R. solanacearum* race 1 biovar 1 (Phylotype III) and race 4 biovar 3 (Phylotype I) are reported as present in locations across Ethiopia (Abdurahman et al., 2017; Lemessa et al., 2010), though race 3 biovar 2 is the dominant genotype in potato production across Ethiopia (Abdurahman et al., 2019) and Sub-Saharan Africa (Abdurahman et al., 2019; Sharma et al., 2022; Sharma et al., 2021). Bacterial wilt caused by race 3 biovar 2, or bacterial wilt henceforth, is particularly difficult to manage due to its ability to survive for over 20 years in a wide range of environmental conditions, hosts, water and soil (Alvarez et al., 2008; Elphinstone, 2005; van Overbeek et al., 2004).

Because potato is generally vegetatively-propagated, seed potato (or potato tubers, as opposed to “true seed”) may have latent bacterial wilt infection, resulting in persistent reinfections and intergenerationally infected seed, threatening Ethiopia’s seed potato production (Abdurahman et al., 2017; Thomas-Sharma et al., 2017; Thomas-Sharma et al., 2016). Methods for managing bacterial wilt once it is present in a field include raising the soil pH, field hygiene and frequently sanitizing tools (Gobena, 2020; Tafesse et al., 2021). More focus is needed on restricting movement of infected seed potato by regulatory bodies, use of certified and Ralstonia-free seed, and establishing an effective monitoring system within the current potato seed trade networks (Carvajal-Yepes et al., 2019; Gobena, 2020; Thomas-Sharma et al., 2016).

Informal potato seed systems are common globally, and potato farmers often save potato tubers from a previous season’s harvest and use these as seed for the next planting season, or sell them to local markets and traders (Abay et al., 2011; Gildemacher et al., 2009). Informal seed trade, with little or no phytosanitary regulation from governmental or institutional actors, provides over 90% of planting material for growers in much of the world and may contribute to the spread of pests and pathogens (Sperling et al., 2020; Sperling and McGuire, 2010; Thomas-Sharma et al., 2016). For this reason, bacterial wilt of potato is becoming a serious concern in East Africa, threatening the rapidly expanding potato industry as well as the food security of the growing population and millions of smallholder farmers.

Formal seed trade networks require collaboration from a wider range of stakeholders, including research institutions and universities, NGOs, policy makers, private seed companies, and regulators (McGuire and Sperling, 2013; Sperling et al., 2020). Formal seed providers allow potato farmers access to disease-free seed, though often the high price, limited availability and low accessibility limit its use; in contrast, informal seed trade provides less expensive and more readily available seed, though these providers often lack information about seed health and quality, potentially resulting in lower yields and infected crops (Hirpa et al., 2010; Yami et al., 2021). Private sector seed businesses are currently underdeveloped in Ethiopia and unable to compete with informal seed prices (Hirpa et al., 2010; Yami et al., 2021). Tools such as the “multi-stakeholder framework” provide an integrated perspective on potato seed stakeholders in formal and informal seed trade networks, highlighting knowledge gaps (Almekinders and Elings, 2001; Bentley et al., 2020; Bentley et al., 2018). A multi-stakeholder approach can identify optimal sectors and locations to take action to meet the needs of each stakeholder group (Andersen Onofre et al., 2021b; Bentley et al., 2018). Actions can include management interventions or implementing certified seed systems to increase phytosanitary standards and international potato seed quality (Choudhury et al., 2017; Garrett, 2021a). The multi-stakeholder framework has been used to identify knowledge gaps in seed systems, and provides a basis for comprehensive analyses which incorporate all the actors in the potato seed chain (Andersen Onofre et al., 2021b; Bentley et al., 2018).

Identifying candidate priority locations for management intervention can help optimize limited resources. Analyzing the networks of trade and potential pest and pathogen movement across cropland production areas can characterize locations at a higher risk for establishment of diseases across Ethiopia, and in countries that are neighbors and/or trading partners (Andersen Onofre et al., 2021b; Buddenhagen et al., 2022; Garcia Figuera et al., 2022; Garrett et al., 2018). Gravity models of epidemic spread can be applied in this context, where areas with a greater abundance of crop hosts, along with a high connectedness to other areas with cropland, may be at a higher risk for pathogen spread and establishment (Jongejans et al., 2015; Rauch, 2016; Xing et al., 2020). Connectedness to other areas is dependent on the type of pest or pathogen movement. For example, the movement of the RSSC in a country is primarily through seed trade. Network analyses where connectivity measures reflect pathogen or pest movement patterns can highlight locations that are likely priorities for management in seed systems.

This study characterized the current formal and informal potato seed networks in Ethiopia, and the geographic risk for the spread and establishment of seedborne potato pathogens and pests, in general and with a focus on bacterial wilt. The objectives of this study were to i) evaluate the current structure of potato stakeholder interactions and communications in the potato seed system, ii) identify locations that are likely to be particularly important for spread of pests and pathogens such as the RSSC, based on potato cropland connectivity, iii) evaluate where management interventions would likely be most efficient to prevent the establishment of bacterial wilt, based on current knowledge, and iv) evaluate the potential spread of bacterial wilt based on formal potato trade with Ethiopia. We synthesized these analyses to support the development of a more robust pest and pathogen surveillance system, and strategies to reduce the spread of infected seed and improve the quality of Ethiopian potato seed systems.

## 2.0 Methods

### 2.1 Stakeholder interactions in Ethiopian potato seed systems

To evaluate the potato seed system in the Ethiopian state Oromia, the multi-stakeholder framework was applied with key stakeholders from across Oromia in September, 2021, asking stakeholders to rank their interactions and communications with each other, using methods as described by Bentley et al. (2018). The 67 stakeholders represented 11 stakeholder categories: research centers and universities (four individuals), policymakers (two), quality-declared seed (QDS) producers (nine), private seed producers (two), ware potato farmers (12), agricultural input dealers (11), potato traders (three), NGO representatives (three), seed regulators (three), cooperative unions (seven), and district and zonal departments of agricultural extension support services (11). Participants were asked to rate their sector interaction with other value chain actors (and vice versa) on a scale of 1-5, where 1 represented no interaction or formal methods of communication between stakeholder sectors, beyond basic print or electronic media, and 5 represented a very strong interaction between stakeholder sectors, including collegial consultations, research, and development activities. The ratings from participants within a category were averaged to estimate the typical interaction and communication between any two stakeholder categories, as perceived by the individual stakeholders.

The average influence reported between each pair of stakeholder categories was assembled in a directed, weighted network. Because the number of participants varied from one stakeholder group to another, we evaluated the relative confidence in the estimate of each interaction level, evaluated as the ratio of the total number of study participants in each stakeholder category and the total number of participants. For example, there were 12 ware potato farmers and two policymakers (a total of 14 participants), where these participants represented the proportion 0.2 of the total number of participants. This proportion was used as a measure of relative confidence, where more stakeholders reporting an interaction gives a higher relative confidence in the interaction level. The adjacency matrix of stakeholder interactions was evaluated and plotted using the igraph (version 1.3.0) package in R (version 4.3.2) (Csardi and Nepusz, 2023; Ihaka and Gentleman, 1996; R Core Team, 2023). Each node in this network represents a stakeholder category, and a link represents the interaction/communication between two categories. The relative confidence level determined for a pair of stakeholders in the survey results was applied as a link weight. We used node strength, the sum of a node’s link weights, as a summary of the overall level of interaction reported for each stakeholder category in the network. The code implementing this analysis, along with all the code and surveys used for this study, is available in GitHub: https://github.com/bereaetherton/SeedSystems_Ethiopia.

### 2.2 Pathogen and pest risk in Ethiopian potato seed systems

We developed an expert knowledge elicitation survey (available in GitHub: https://github.com/bereaetherton/SeedSystems_Ethiopia) to capture the current domain knowledge regarding Ethiopian potato seed systems and implemented it using Google Forms (Ford and Sterman, 1998). The questions focused on current potato management strategies, pest and pathogen risk, and observed trade patterns. Twenty experts on potato health in Ethiopia participated in the expert knowledge elicitation workshop in Addis Ababa, Ethiopia, on March 13^th^, 2023. The workshop began by characterizing the experts and their domain knowledge across Ethiopia (Figure 1), and then by characterizing the potato growing population and demographics. Next, the experts characterized the observed seed trade networks and the types of seed available to growers. Finally, experts elaborated on parameters specific to bacterial wilt of potato, including the social networks for potato growers who share information about managing bacterial wilt (discussed more in 2.5). Responses from the experts were compiled and represented using the ggplot2 (version 3.3.6) (Wickham, 2016) and rgeoboundaries (version 0.0.0) (Dicko, 2023) packages in R. Some of the results from this expert knowledge elicitation are reported as part of a larger synthesis of banana, cassava, potato, and sweetpotato crops in Cameroon and Ethiopia in Mouafo-Tchinda *et al*. (in review). The main use of the expert knowledge elicitation results in the study reported here is specific to bacterial wilt scenario analyses, while Mouafo-Tchina *et al*. (in review) describe pests and pathogens and their role in humanitarian contexts.

**Figure 1.**
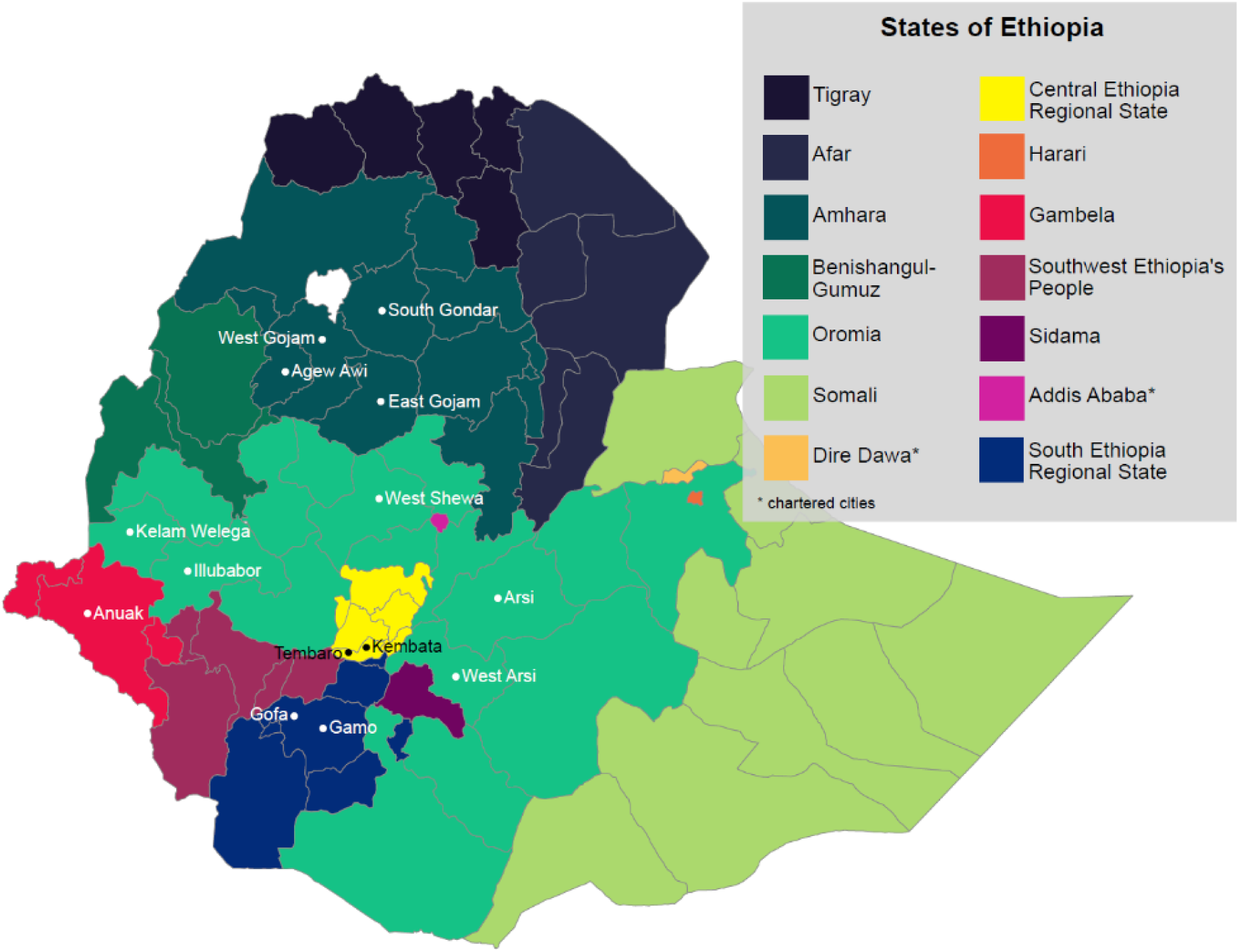
The states and chartered cities of Ethiopia, along with several zones referenced in the text. Zonal administrative boundaries reflect 2023 zonal administrations, and the point next to a zone name indicates the approximate centroid of the zone.

### 2.3 The Ethiopian potato cropland network: key roles of locations for surveillance

The likely importance of a node (in this case, a location with potato production) in an epidemic or invasion network can be evaluated in several ways. Here we apply, first, a risk index based on cropland connectivity, and later, as described below, epidemic simulations and scenario analyses. For each of these analyses, the landscape of potato production in Ethiopia was modeled as a potential dispersal network for potato-specific pathogens and pests, based on potato cropland distribution data in MapSPAM (IFPRI, 2020). Raster cell values represented the potato harvested area (ha) in each cell for 2017, the most recent estimates available. The cell values were divided by 10,000 (equivalent to m^2^/ha) to obtain the proportion of harvested area, and the resolution was adjusted by aggregating from 5-min to 10-min (Arc) by calculating cell means (merging a square of four adjacent cells) for simplicity. The potato cropland data for Ethiopia and the surrounding area was selected. Cells with a proportion of harvested potato area greater than 0 were kept in the analysis. Host availability was represented using a gravity model (Jongejans et al., 2015), using methods described by Andersen Onofre et al. (2021b) and Xing et al. (2020). The entries in the adjacency matrix, *M*_*ij*_, are the relative likelihood of pathogen movement between the locations (nodes) i and j, based on a negative exponential function (*M*_*ij*_ = *ac*_*i*_*e*_*j*_*e*^−*ydij*^, where *γ* is the dispersal parameter) or an inverse power law model (*M*_*ij*_ = *ac e d*^−*β*^) where *e* and *e* are the proportion of potato harvested area for nodes i and j, *d* is the distance between the two nodes, *β* is a dispersal parameter, and *α* is a scaling parameter (Rauch, 2016). The probability of movement between two nodes decreases with distance as a function of β or *γ*; for example, the likelihood of long-distance movement when β = 0.1 is much greater than when β = 0.5; similarly, the likelihood of movement is much greater for *d*_*ij*_ = 1 than *d*_*ij*_ = 10. The geographic coordinate of each cell was extracted using function extentFromCells in the terra package (version 3.5.15) in R (Hijmans et al., 2022), and the Euclidian distance between each pair of nodes was calculated.

In sensitivity analyses, we evaluated the responses of a cropland connectivity risk index (CCRI) to a range of potential dispersal gradients in the gravity model, as in Xing et al. (2020). The parameter values β = 0.5, 1, and 1.5 were used for the inverse power law function and *γ* = 0.05, 0.1, and 0.2 for the negative exponential function. The cropland density matrix and distance matrix were rescaled between 0 and 1, and then multiplied element-wise. For perspective, two states each with a cropland density of 1 ha, that are 10 km apart, would have a relative likelihood of pathogen movement equal to 0.31, 0.1, and 0.03 for β = 0.5, 1, and 1.5, respectively. If cropland density were to increase, or the distance between two nodes were to decrease, the relative likelihood of movement would increase.

We generated maps of this CCRI in R for each network generated in the sensitivity analysis, as in Xing et al. (2020). Higher CCRI indicates locations that are more likely to play a significant role in epidemics or invasions, based on host availability, and thus are candidate priorities for monitoring. This CCRI was defined as the weighted sum of four network metrics, to capture a range of potential roles of a node in invasion networks: 1/2 the node betweenness centrality to account for the role of a node in shortest pathways, 1/6 node strength to represent the role of a node’s neighbors, 1/6 a node’s nearest neighbors’ node degrees to account for the role of neighbors’ neighbors, and 1/6 eigenvector centrality to account for the node’s position in the larger network. The relationship between crop density and the CCRI scores was linear in these analyses. The CCRI tends to be higher where there is more potato production, but also is relatively higher than potato density for locations with high proximity to many other potato production locations, and for locations with high betweenness centrality, that may act as a bridge between potato production areas. The potential for a pathogen to spread through the crop landscape was evaluated for the general dispersal of pests and pathogens in the potato landscape, including *R. solanacearum* race 3, in the sensitivity analysis.

Cropland connectivity analysis functions are available in the geohabnet R package (<u>cran.r-</u> <u>project.org/package=geohabnet</u>). This analysis and others are part of the open-source R2M toolbox for rapid risk assessment to support mitigation of pathogens and pests (www.garrettlab.com/r2m), with example applications including Andersen et al. (2019), Andersen Onofre et al. (2021a), Buddenhagen et al. (2017), Etherton et al. (2023), and Nduwimana et al. (2022).

### 2.4 Candidate surveillance locations based on epidemic simulations

While the analysis of CCRI above is based on standard node centrality measures, an alternative approach to evaluating the potential role of locations in surveillance is based on epidemic simulation. We simulated epidemics in Ethiopia’s potato landscape, based on the invasion network structure described above, using the smartsurv function in the impact network analysis (INA) package (version 1.0.0) (Garrett, 2021b) in R. This function evaluates how many other locations would likely still be free of an invasive pathogen or pest when the invasion is detected at a particular surveillance location. It considers each location in turn as a potential monitoring site. We evaluated this measure of efficacy for surveillance for each location in Ethiopia with potato production, in sensitivity analyses based on approximately the same adjacency matrices as described in 2.3, for parameter values β = 0.5, 1, or 1.5 and *γ* = 0.05, 0.1, or 0.2. These scenarios were based on each node (location) having an equal likelihood of being the point of initial pest or pathogen invasion. The analyses were repeated in 400 realizations for each parameter combination. The means of the results in the sensitivity analysis of the smartsurv score were rescaled between 0 and 1 and taken as measures of a location’s efficiency for detecting pathogen or pest invasion. The relationship between smartsurv scores and cropland density was exponential in this analysis, where the smartsurv score increased rapidly for cropland densities between 0 and 50 ha and increased slowly from 50 to above 300 ha per pixel. This suggests that the results of the epidemic simulations will highlight locations with sufficient host density to play an important role in epidemic networks.

### 2.5 Potential management interventions and impact network analysis

A more detailed scenario analysis of potential pathogen establishment was evaluated in potato cropland pixels across Ethiopia using the INAscene function in the INA package (version 1.0.0) (Garrett, 2021b) in R. This scenario analysis incorporated both (a) soil pH as an environmental factor influencing bacterial wilt risk (Li et al., 2017; Nion and Toyota, 2015; Tafesse et al., 2021) and (b) an associated social network of management decision-makers linked to the potential invasion network. Tafesse et al. (2021) estimated bacterial wilt incidence as a function of soil pH, indicating that bacterial wilt incidence for strains collected in Ethiopia is less likely above a pH of 6.5. Similarly Li et al. (2017) found that the range of pH conducive to *R. solanacearum* growth was between 4.5 and 5.5 in a field setting. We used the 2012 FAO topsoil pH maps to evaluate the relative likelihood of bacterial wilt establishment based on pH (FAO, 2012). Pixels with a pH 4.5-5.5 were given weight 1.0, pixels with pH 5.5-6.5 were assigned weight 0.5, and all other pixels were assigned weight 0.1. Cropland data from MapSPAM (IFPRI, 2020) potato cropland distribution data were aggregated to 15-min by 15-min (Arc) for simplicity using the terra package in R (Hijmans et al., 2022).

The INAscene function simulates a multilayer social-ecological network system with an adjacency matrix of weights used to simulate the spread of a pathogen in an invasion network, and the associated network of communication that influences decision making, where the potential spread of a pathogen and spread of information and influence occur in each time step (Garrett, 2021b). Here, the results from the expert knowledge elicitation (described in 2.2) were used to create a social adjacency matrix and a pathogen invasion adjacency matrix. Experts were asked to identify potato seed trade and farmer communication networks which occurred within a state and between states. These networks were averaged across experts and rescaled between 0 and 1. The initial locations of disease in scenario analyses were selected as the locations with conducive pH (between 4.5-5.5). Scenarios were evaluated using INAscene for 10 timesteps in 400 realizations for each of a range of management adoption probabilities. The rate of bacterial wilt establishment at a node was averaged across the 400 realizations for each scenario (parameter combination) and used to determine locations at high risk for *R. solanacearum* establishment.

We also evaluated the combined results of these three approaches for identifying key locations based on cropland networks: a cropland connectivity risk index (based on epidemic network centrality metrics), simulated epidemics (smartsurv output), and scenario analyses incorporating environmental conduciveness based on pH and social networks influencing management (INAscene output). These results were combined in a visualization with MapSPAM potato cropland density and the FAO pH maps using the biscale package (version 1.0.0) (Prener et al., 2022) in R. Crop density and pH were used to create mapping classes for a bivariate map, and a kernel density estimation was performed using the results of cropland connectivity analyses, epidemic simulation, and scenario analysis.

### 2.6 International potato seed trade and invasion risk networks

The network of countries formally trading potato directly with Ethiopia was characterized based on the annual export and import quantity of fresh potatoes (FAOSTAT, 2020). Potato flour, potato offal, and frozen potato were excluded from the analyses as these products are processed and unlikely to spread pathogens or field pests. Annual mean trade activity between each pair of reporter-partner countries from the 2005-2019 reports of exports and imports was calculated. The most recent reported geographic distribution of bacterial wilt was generated from recent reports of the Centre for Agriculture and Bioscience International database (CABI, 2020) combined with the EPPO dataset (EPPO, 2021). The pathogen invasion potential for a country was calculated based on the log-transformed potato harvested area in hectares (FAOSTAT (2020)), and the reported status for bacterial wilt of potato in each country (CABI, 2020). The pathogen trade movement potential between countries was determined using the gravity model *M*_*ij*_ = *e*_*i*_*e*_*j*_*t*_*ij*_, where *e*_*i*_and *e*_*j*_represent the pathogen invasion potential of countries i and j, and *t*_*ij*_is the log-transformed potato trade (mt) between country pairs. Trade network visualizations, in which nodes represent countries, were built with weighted links indicating the pathogen trade movement potential between countries (*M*_*ij*_).

Potato trade networks were examined for Ethiopia in two ways: 1) including only the direct trading neighbors and 2) including other partners two steps away from Ethiopia. The potential epidemic risk to a country is summarized by calculating its node in-strength, that is, the sum of link weights associated with commodity imports (Brooks et al., 2008; Bucur and Holme, 2020; Silk et al., 2017).

## 3.0 Results

### 3.1 Multi-stakeholder interactions in potato seed systems

The multi-stakeholder interaction network reported by stakeholders (Figure 2, A) indicates several stakeholder sectors with high levels of communication and interactions: research centers and universities with policymakers, NGOs and seed regulators; quality declared seed (QDS) producers with NGOs; cooperative unions with NGOs; ware potato farmers with potato traders; and seed regulators with policy makers (Figure 2, D). These high reported interactions, however, all had lower relative confidence (< 0.25) given the lower representation of individuals from some stakeholder sectors in the study. Low levels of communication and interaction were reported for the following: ware potato farmers’ relationship with policymakers; NGOs and seed regulators with ware potato farmers; and research centers, policymakers, seed regulators, seed companies and NGOs relationship with potato traders (Figure 2, B). The ware potato farmers had the highest relative confidence scores (> 0.25), with extension support and then input dealers also having higher relative confidence scores, given their higher representation in the study compared to other stakeholders (Figure 2, C).

**Figure 2.**
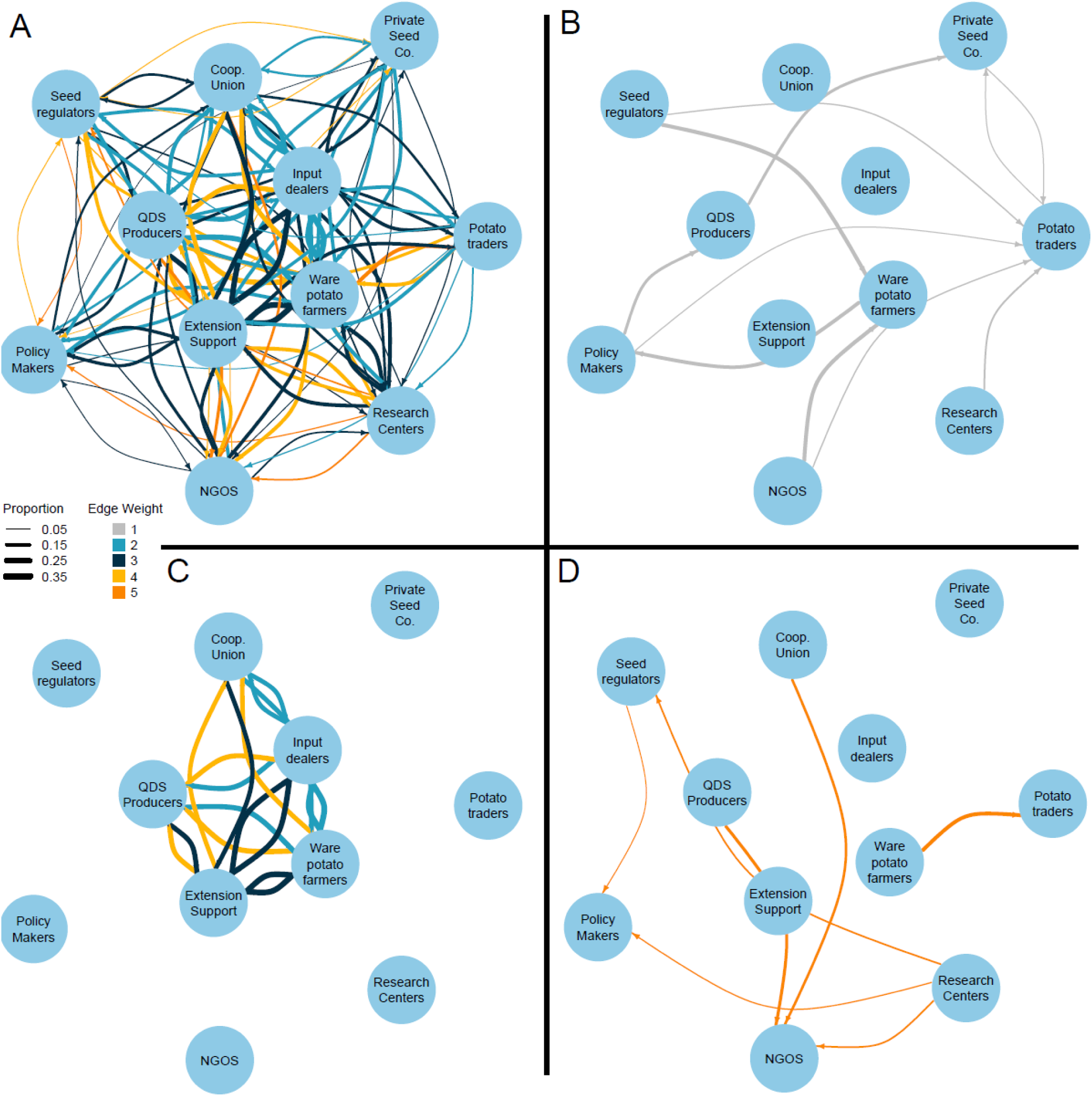
Reported networks of interactions between potato system stakeholders in Oromia state, Ethiopia. Reported interactions ranged from 1, no interaction, to 5, high interaction. Nodes represent stakeholder groups, link color indicates the average reported interaction levels, and link width is proportional to the relative confidence level for interactions, based on the proportion of individuals representing each stakeholder group. A) The complete stakeholder interaction network, with reported no-interaction links excluded for simplicity. B) The stakeholder network indicating only reported no-interaction links, which are potential targets for system improvement. C) The network with only interactions reported as greater than 1 and with relative confidence greater than 0.25. D) The network with only the reported high interactions.

Research centers and universities had the highest reported levels of interaction and communication across all stakeholder sectors, with a total node strength of 65, the sum of ratings (node strengths) for stakeholders were for NGOs (64), seed regulators (61), agricultural extension services (61) and QDS producers (61). Potato traders had the lowest reported node strength (45) across the stakeholder sectors. Input dealers had the next lowest node strength (49).

### 3.2 Geographic priorities indicated by cropland connectivity, epidemic simulations, and INA scenario analysis

In the cropland connectivity analyses, there were eight locations (nodes) with a higher CCRI score of approximately 0.7 (Figure 3, A), in Gamo, Gofa and central South Gondar. There were large areas of cropland with a CCRI greater than 0.4 in Kembata, Tembaro, Arsi, West Arsi, Agew Awi, West Shewa, and East Gojam. States with a higher CCRI score typically had high cropland density, and were located in the states of Oromia, Amhara or the Southern Nations Regional States. For surveillance within a state, locations with high CCRI scores could be candidate priorities for surveillance.

**Figure 3.**
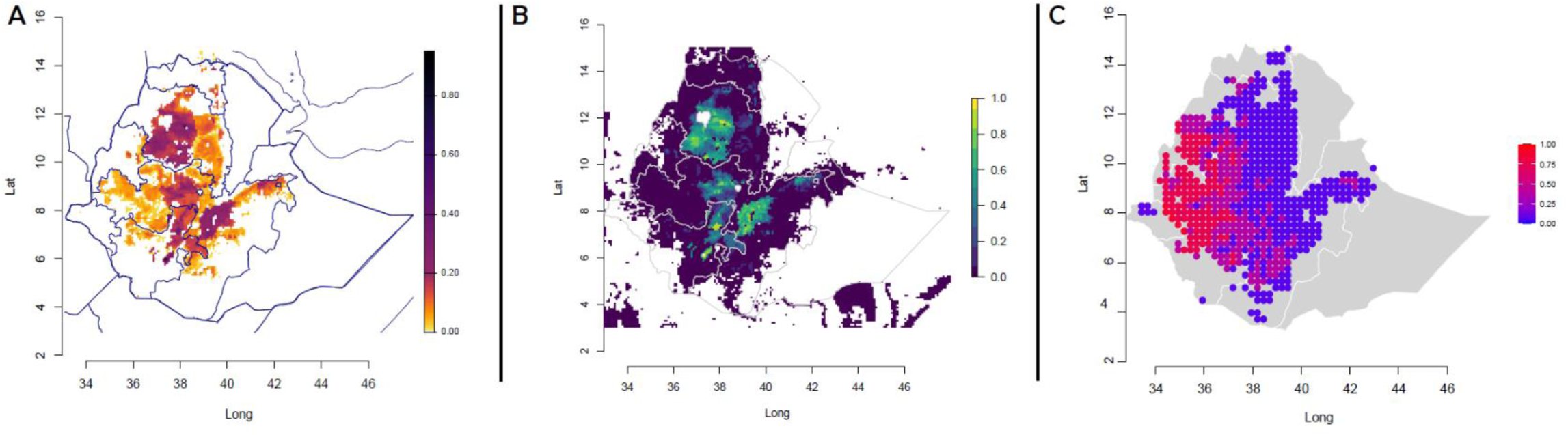
Three perspectives on the importance of locations in Ethiopia for surveillance and mitigation of potato pathogens and pests. A) Candidate priority locations based on epidemic simulations (smartsurv analyses) are indicated by brighter colors. B) Candidates based on a cropland connectivity risk index (CCRI) are indicated by darker colors. C) Candidates based on social-ecological scenario analyses (INAscene analyses) are indicated by brighter colors, where the scenario analysis incorporates factors important for bacterial wilt establishment, including soil pH suitability. These perspectives are integrated in Figure 4.

The epidemic simulations (using the smartsurv function) indicated how far a pathogen would likely spread in the invasion network before detection at a given location. That is, for each location (node) taken as a focus, the output is the mean proportion of other locations in the network that remained free of the pathogen at the time when the pathogen was detected at that focus node. The more locations that remain pathogen-free when the pathogen is detected at the focus location, the more efficient the focus location is for surveillance in the simulations. Based on this analysis, candidate locations for surveillance at the national level included potato production locations in zones Gamo, Gofa, South Gondar, Kembata and Tembaro, with proportions greater than 0.9. If bacterial wilt was detected in these locations in the simulations, this information would trigger mitigation before it moved to 800 other locations, where each location in this analysis represents roughly 8,600 ha of potato cropland (Figure 3, B). Other candidate locations for surveillance, or locations with a smartsurv score greater than 0.7, included cropland locations in zones Arsi, West Arsi, West Shewa, Agew Awi, West Gojam and South Gondar. As in the CCRI results, at a national level these locations were primarily found in Oromia, Amhara and the Southern Nations Regional States (Figure 3, B), and candidates for surveillance within a state would include locations with locally high values.

The scenario analyses (using the INAscene function) included the potential effects of soil pH on disease and system parameters provided by expert knowledge elicitation. In these simulations, bacterial wilt establishment was likely to occur along the border between the zones Anuak, Illubabor and Kelam Welega in more than 80% of realizations (Figure 3, C). Bacterial wilt establishment occurred across eastern Ethiopia, in Oromia, Benishangul-Gumuz, Amhara and the Southern Nations, in more than 70% of all realizations. All these locations were reported to have pH below 6, and a moderately high potato crop density. The CCRI, epidemic simulations, and scenario analyses, when combined into a biscale map of crop density and pH, identified candidate priority locations for surveillance in Agew Awi and in the area between Kembata, Tembaro, Gamo and Gofa (Figure 4). These locations could have important roles in bacterial wilt invasion given their CCRI, were identified as candidate locations for surveillance given their smartsurv scores, and had higher bacterial wilt establishment in INAscene scenarios. These zones all have relatively high cropland density and a low soil pH, and are identified as candidate priority locations for national surveillance of potato bacterial wilt. Furthermore, South Gondar and southern Arsi have candidate priority locations for surveillance based on the CCRI and smartsurv results, which did not incorporate pH as a factor. In these areas the pH is not likely to be as conducive for bacterial wilt establishment compared to other areas, but these locations may be priorities for other pathogens and pests.

**Figure 4.**
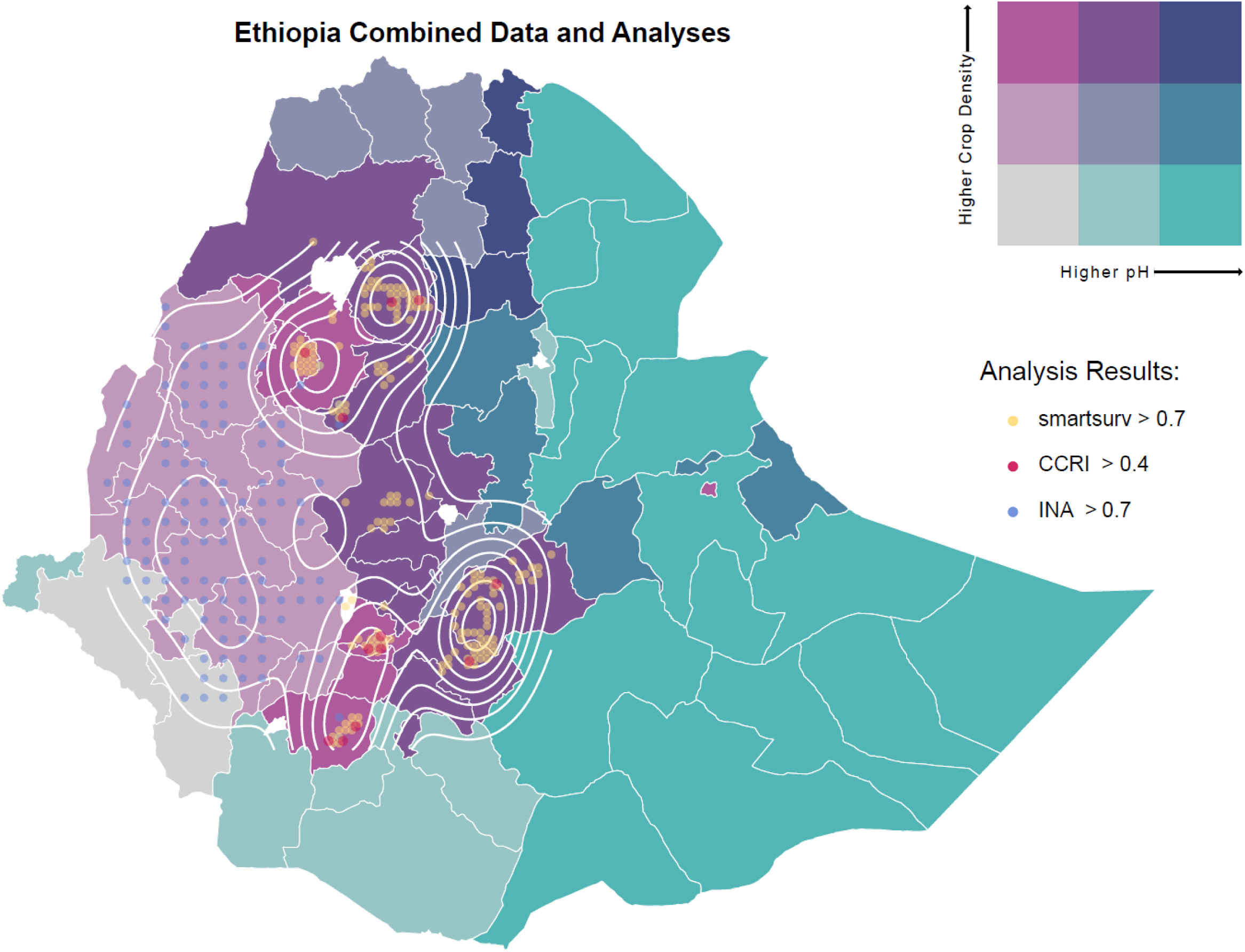
Candidate priority locations for management of potato pathogens and pests in Ethiopia. The results of three analyses illustrated in Figure 3 are combined here: cropland connectivity (CCRI), epidemic simulation (using the smartsurv function) and scenario analyses (using the INAscene function). The scenario analysis is more specific to the risk of bacterial wilt, incorporating pH as an epidemiological risk factor. Points show locations that scored highest in these analyses and are layered on a biscale map, where color represents the relative pH and potato density. A density layer was added to show where the highest frequency of points are located. Zones that are a focus of the results are labeled in detail in Figure 1.

### 3.3 International potato seed trade with Ethiopia

To assess the potential for introduction of new pathogens and pests to Ethiopia via international potato imports, and for re-introduction of *R. solanacearum* race 3, we analyzed the Ethiopian trade network of formal potato imports and exports in 2005-2019. Ethiopia imported fresh potatoes from 12 countries, of which six, including China, France, Germany, India, Kenya and the Netherlands, have reported cases of *R. solanacearum* (Figure 5, A). During the same period, Ethiopia exported fresh potatoes to seven countries including Djibouti, Russia, Saudi Arabia (SB), Sudan, former Sudan, and the United Arab Emirates (UAE), while *R. solanacearum* has not been reported in these countries.

**Figure 5.**
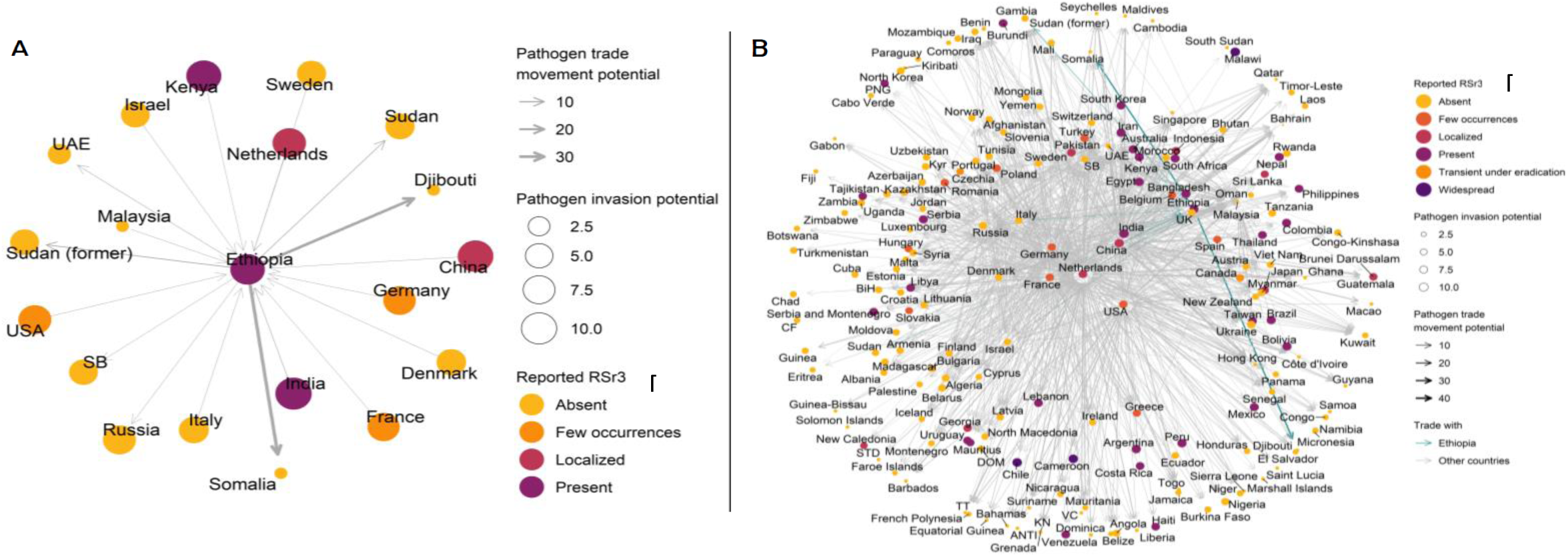
Formal trade networks reported for potato (A) one step from Ethiopia and (B) two steps from Ethiopia (data: FAO). Link size represents the invasion potential (or the relative trade amount in metric tons), the node size represents a measure of the potential for pathogen invasion (based on the trade amount, reported pest status and reported harvested cropland), and the color represents the reported presence of *Ralstonia solanaceaum* race 3 (data: CABI).

The presence of hubs in global trade networks is another important aspect to consider when analyzing the potential movement of pathogens associated with international trade. In the network analysis (Figure 5, B), international trade hubs were characterized by high levels of both importation and exportation, and these had the highest node strength in the trade network. By using a network layout that positions these hub countries toward the center of the network, this analysis highlights the potential role of international trade hubs for pathogen movement for three cases. 1) Pathogen introduction is a risk for countries where the pathogen is not present if they import from international trade hubs where the pathogen is present. 2) International trade hubs may be important if they re-export commodities from countries where the pathogen has invaded to countries where the pathogen is absent. 3) Pathogen re-introduction is a risk for international trade hubs if importing commodities from countries where a pathogen is present. The analysis indicates that Ethiopia has strong trade connections with important international trade hubs including countries exporting to Ethiopia (Netherlands, India, China) and importing from Ethiopia (Russia and France; Figure 5, B).

## 4.0 Discussion

In the absence of a formal seed certification scheme and regional quarantine measures in Ethiopia, bacterial wilt is a serious concern, threatening the potato industry. The multi-stakeholder analysis of Ethiopia’s potato seed system, highlighting Oromia, indicated weak direct interactions between ware potato farmers and other actors in the potato value chain (Figure 2, B) with the exception of potato traders. A majority of the smallholder ware potato farmers in Ethiopia participate in internal informal trade of seed potato with high risk of disease dissemination to potential uninfected areas. Farmers often obtain knowledge regarding seed quality and management practices informally as well. One step toward ensuring the food and income security of smallholders in the potato farming systems could be greater involvement of key system stakeholders to support the quality and health of seed potato (Beyene, 2010; Carvajal-Yepes et al., 2019; Damtew et al., 2018; Haverkort et al., 2012).

The interactions between the ware potato farmers and research centers, public extension and NGOs may provide limited information regarding disease management, and despite current efforts to build a formal system, informal farmer-based information and seed sources continue to prevail across most states in Ethiopia (Almekinders et al., 2019a; Almekinders et al., 2019b; Bèye and Wopereis, 2014; Damtew et al., 2018). A shift to a more preventative management culture warrants networking of local level actors who can collectively engage in disease monitoring and information sharing on key aspects of bacterial wilt management. Agricultural stakeholders need to consider potato farmers’ socioeconomic needs and focus on strengthening the growers’ knowledge networks for disease management strategies (Cravey et al., 2001; Sligo and Massey, 2007).

When farmers have little access to educational resources and management information regarding pathogen spread, the potential for disease expansion increases, particularly when producers’ understanding of crop management is in conflict with the recommended best practices (Etherton et al., 2023; Šūmane et al., 2018). For example, Gemeda et al. (2020) found that most pastoralists across Ethiopia treat their livestock with antibiotics, though a high proportion did so incorrectly by not following the full antibiotic treatment courses, potentially increasing the risk of resistant microbial pathogens in livestock. Okello et al. (2020) found that a majority of potato farmers in Kenya were unable to correctly diagnose bacterial wilt, and Uwamahoro et al. (2018) found that only about 50% of farmers in Rwanda were aware of the major dispersal mechanisms for bacterial wilt.

The multi-stakeholder analysis found a large gap in the relationships in the potato value chain. This suggests there is an opportunity to increase the quality of seed potato in Ethiopia by incorporating all actors in the potato seed chain through education and building disease management knowledge (Andersen Onofre et al., 2021b; Bentley et al., 2018; Damtew et al., 2018). Strengthening the stakeholder relationships could help establish a seed trade system that integrates both the formal and informal seed trade networks, in which policy makers, input dealers, and private seed companies can take the lead in seed potato certification processes and quality seed delivery to more ware potato farmers (Almekinders and Elings, 2001; Bentley et al., 2018; Choudhury et al., 2017; Garrett, 2021a; Thomas-Sharma et al., 2017).

Increasing communication among stakeholders can help to prevent the spread of pests and pathogens, but it is also important to consider actively monitoring and surveilling potato cropland for bacterial wilt. The cropland connectivity analyses found locations in two primary states, in Gamo, Gofa and central south Gondar, that are likely to be particularly important for epidemic spread through the national potato production landscape. Given their higher cropland connectivity, these are candidate locations for regular sampling for potato pathogens and pests. Analyses in epidemic simulations using smartsurv found similar results, with locations in Gamo, Gofa, South Gondar, Kembata and Tembaro as candidates for potato pest and pathogen surveillance. These locations are likely to be efficient for national surveillance, with a higher likelihood for detection of early disease expansion, allowing more time for action in other locations before a pathogen or pest spreads widely.

The cropland connectivity (CCRI) and epidemic simulation (smartsurv) analyses were not applied here using disease-specific parameters, and so were designed for application to a range of potato pests and pathogens. An expert knowledge elicitation workshop identified late blight, potato virus Y, early blight and potato leafroll virus as threats to potato production in Ethiopia, consistent with previous studies (Guchi, 2015; Tafesse et al., 2018). The scenario analysis using INAscene was specific to bacterial wilt, designed based on current knowledge about bacterial wilt and with the results from expert knowledge elicitation (Ayana et al., 2011; Li et al., 2017; Tafesse et al., 2021; Tafesse et al., 2020; Tessema and Seid, 2023). The scenario analyses, which included the potential effects of pH on bacterial wilt establishment, found that disease infection and spread were more likely to occur in locations along the border between Anuak, Illubabor and Kelam Welega. These locations have a lower potato cropland density, but higher potential for disease establishment due to pH effects. However, the overall magnitude of the economic impact due to disease establishment in these locations may be lower than in areas with more intensive potato production.

Epidemics or invasions generally occur in phases, starting with the introduction of a pathogen or pest, continuing with expansion and establishment, and potentially ending with the extinction of susceptible hosts (Grenfell and Harwood, 1997; Hui and Richardson, 2017).

Strategies for managing and detecting disease differ greatly across these phases; for example, if disease is managed effectively during the lag (or introductory) phase of an epidemic, disease eradication may be feasible, as discussed in Etherton et al. (2023). Results from the cropland connectivity, epidemic simulation, and scenario analyses identified locations that are important candidates for pathogen surveillance, though the value of these locations depends on the phase of the epidemic. Bacterial wilt is now reported across Ethiopia, with disease incidence up to 63% on potato (Kurabachew and Ayana, 2017; Tessema and Seid, 2023). The expert knowledge elicitation indicated bacterial wilt presence in all states of Ethiopia. For zones where bacterial wilt is not yet present, the scenario analyses may help identify locations at high risk for disease establishment and priorities for disease management. For zones where bacterial wilt incidence is still low, the epidemic simulation results may be well suited, as these results are targeted to stop spread and monitor for bacterial wilt. For zones where bacterial wilt incidence is high, candidate management priorities are locations with a higher CCRI score, as these locations are highly connected and have the potential to contribute to infected seed trade. These results are particularly important for policymakers, regulatory bodies and researchers to develop more effective prevention and disease management strategies at local, regional and national levels.

Based on the analysis of international potato trade, Ethiopia may be at risk for reintroduction of the RSSC from potato seed imported from India, Kenya, China, and the Netherlands, unless seed is delivered in confirmed aseptic form. Seed potato imported from these areas should be thoroughly tested for the absence of the RSSC using reliable diagnostic tools, since these countries have reported the presence of bacterial wilt. Equally, seed potato exported from Ethiopia should be tested for the absence of the RSSC, particularly to countries where bacterial wilt has not yet been reported, such as Denmark, the Netherlands, and Italy. Ethiopia also exports potato to Southern Sudan, Sudan, Djibouti, and Somalia, where bacterial wilt has not yet been reported. An important question for interpreting these trade structures and their associated risk is whether bacterial wilt is currently absent or rare in Sudan, Djibouti, and Somalia, or has simply not been reported yet. If it is currently absent or rare, consideration of the risk of bacterial wilt spread through potato trade is important, since bacterial wilt is persistent for long periods of time in soil, water and environment (Ayana et al., 2011). However, civil unrest in some of these countries may make extensive monitoring of potato trade impractical in the short run. It also may increase the risk of disease introduction when there is need for rapid movement of food aid (Mouafo-Tchinda *et al.,* in review; Etherton *et al*., in review).

There is no single management option for the complete eradication of bacterial wilt, however, growers can sanitize tools, practice crop rotation or intercropping, and regularly replace informal potato seed with certified seed (Okello et al., 2020; Uwamahoro et al., 2018). Regional adoption of these management strategies and the active monitoring of potato seed quality and movement are complicated by many factors, for example, land contraints, crop intensification and soil nutrient depletion (Headey and Jayne, 2014; McGuire and Sperling, 2013). Adoption of any management strategy primarily depends on its utility and farmers’ ability to access it in an increasingly complex future (Almekinders et al., 2019b; Carvajal-Yepes et al., 2019; Thomas-Sharma et al., 2017). In addition to bacterial wilt, the challenges posed by other pulse and press stressors, in the context of disaster plant pathology (Etherton *et al.,* in review), play a role in management adoption and the success of formal seed systems. For example, recent pulse stressors like the COVID-19 pandemic and the Ethiopian civil war crisis cause mass migration as well as a decline in available clean seed, whereas press stressors like climate change, mixed patterns of rainfall, and droughts, all potentially restrict and disrupt consistent monitoring and testing for bacterial wilt and other pathogens (Conway and Schipper, 2011; Longley et al., 2002).

The spread and establishment of major pests and pathogens through seed trade networks can be mitigated by increased involvement of stakeholders and effective monitoring of high-risk areas for spread and establishment. This assessment evaluated the existing interactions between stakeholders and identified wide gaps among value chain actors; closing these gaps could increase potato farmer knowledge and reduce the spread of the RSSC in Ethiopia. We identified areas across Ethiopia which may be at high risk for potato pest and pathogen establishment and spread, with a focus on scenario analyses for bacterial wilt. Disease surveillance and seed testing need to be strengthened, with important candidate locations in Agew Awi, and the stretch between Kembata, Tembaro, Gamo, and Gofa, as surveillance in these areas is likely to make efficient use of seed testing resources given their position in Ethiopia’s potato seed system. Lastly, countries that exchange seed with Ethiopia should be cautious of bacterial wilt spread, and Ethiopia should regularly test seed potato imported from other countries with reported bacterial wilt presence. This study can support the informed intervention of potato value chain stakeholders, particularly policymakers, for the mitigation of pathogen and pest spread, regionally and internationally.

## Acknowledgements

This research was undertaken as part of, and funded by, the CGIAR Seed Equal Research Program, and by the CGIAR Research Program on Roots, Tubers and Bananas (RTB), and supported by CGIAR Trust Fund contributors: https://www.cgiar.org/funders/. We appreciate helpful input from Setegn Gebeyehu and Mihiretu Cherinet.

## Contributions

B. A. Etherton, J. Yao, A. I. Plex Sulá, V. S. Kosmakos, R. A. Mouafo-Tchinda, K. Sharma, and K. A. Garrett developed the initial manuscript drafts. F. Asfaw, R. Kakuhenzire, H. A. Kassaye, R. A. Mouafo-Tchinda, and K. A. Garrett performed field surveys and stakeholder surveys and implemented expert knowledge elicitation for Ethiopian seed systems. B. A. Etherton, A. I. Plex Sulá, R. W. McCoy, Y. Xing, and K. A. Garrett developed and implemented data analyses. All authors reviewed and contributed to the final manuscript.

